# “Shedding light on plant proteolysis: genetically encoded fluorescent sensors as tools for profiling protease activities.”

**DOI:** 10.1101/2024.06.08.598063

**Authors:** Álvaro Daniel Fernández-Fernández, Simon Tack, Matthias Van Durme, Jonah Nolf, Moritz K. Nowack, Jens Staal, Simon Stael, Frank Van Breusegem

## Abstract

Proteolysis, a ubiquitous process in living organisms, is driven by proteases that regulate numerous signaling pathways through the hydrolysis of peptide bonds in protein substrates. Understanding the temporal and spatial dynamics of proteolysis and the activation of proteases is crucial for elucidating their roles in biological pathways. Here, we introduce a suite of genetically encoded FRET reporters designed to detect various proteolytic activities in plants. These sensors effectively reported *in planta* the specific activity of both Tobacco Etch Virus protease and caspase-3. Furthermore, we developed sensors for detecting plant metacaspase activity, validated through both *in vitro* and *in planta* experiments. These experiments revealed the spatial dynamics of proteolysis triggered by metacaspase activation following wounding and programmed cell death in roots. The implementation of these tools in plant biology research opens new avenues for investigating proteolytic mechanisms, significantly enhancing the potential for in-depth studies. Our work demonstrates the feasibility of using these sensors to detect diverse protease activities *in vivo* with high spatiotemporal resolution. These plant proteolytic biosensors hence represent a valuable toolbox for understanding protease functions within their natural context, paving the way for future advancements in plant biology research.

## INTRODUCTION

Protein proteolysis is a fundamental and crucial aspect of every biological system ^1,2^, ensuring the removal of damaged or unnecessary proteins, and contributing to the maintenance of cellular health and functionality ^3^. But proteolysis is not merely a protein breakdown process; rather, it serves also as a vital regulatory mechanism essential for the dynamic modulation of cellular processes in plants and animals^4,5^. Protein hydrolysis generates novel protein variants with distinct properties and lifespans. Through proteolysis, multiple cellular processes are controlled, nutrient recycling is facilitated, and responsive adjustments to environmental changes are enabled. Peptidyl hydrolysis is carried out by proteases, enzymes that break peptide bonds within proteins. Consequently, proteolytic activity needs to be meticulously regulated, with many proteases initially synthesized as inactive precursors known as zymogens. Understanding proteases within their native environments and the mechanisms underlying their activation is fundamental for elucidating their functions and contributions to particular physiological pathways.

In plants, the number of genes encoding for proteases is notable. For instance, in species like *Arabidopsis thaliana* and *Populus trichocarpa* between 600 to 800 proteases are annotated ^4,6,7^. Plant proteases are reported to play integral roles in diverse cellular events, ranging from organellar protein import ^8^ to control of growth and development ^9,10^. Additionally, proteases contribute to plant adaptation, responses to abiotic stresses, immunity, hypersensitive response, and developmental programmed cell death ^5,11–13^. Despite the high number of protease families and their relevance in numerous cellular processes, the specific mode of action and activation mechanisms of plant proteases remain largely undiscovered with a few exceptions ^14,15^. Among plant proteases, the metacaspase family of cysteine proteases has garnered significant attention in the past decade. Metacaspases play critical roles a.o. in programmed cell death and defense responses ^16–25^. The activation mechanisms of most plant metacaspases rely on elevated calcium levels, prompting self-processing and subsequent activation^19,26,27^, except for a subgroup of pH-controlled metacaspases^28–30^. In Arabidopsis, calcium-activated metacaspases require millimolar levels of this cation for their activation. During wound stresses, calcium levels increase in the local area next to the damaged tissue triggering metacaspases self-processing and subsequent cleavage of their substrates such as PROPEPs. In Arabidopsis roots, AtMC4 can process AtPROPEP1, which resides in the tonoplast layer, releasing its mature form AtPep1^19^. AtPep1 and other members of the Pep family function as damage associated molecular patterns (DAMPs). After maturation, they bind to the receptors, PEPR1 and its homologue PEPR2, initiating a cascade of defense responses^31^.

Previously, in mammalian cell lines, proteolytic activities were visualized and quantified *in situ* using genetically encoded reporters ^32–39^. While genetically biosensors monitoring redox or metabolite homeostasis are already successfully implemented at organ and cellular level in plant biology^40,41^, tools for the *in vivo* assessment of proteolytic activity in plants were limited to fluorogenic activity-based protein profiling probes^39,42^. Notable exceptions in plants are BRET sensors, used to detect cysteine protease ATG4 activity in autophagy ^43^ and to FRET sensors for real time detection of DEVDase activity during UV-stress^44^. Despite their potential, biosensors remain underutilized in the study of plant proteases. Developing advanced biosensor tools to monitor proteolytic activities will significantly advance fundamental plant science and might inspire for practical applications for improving crop productivity and sustainability.

Here, we developed a highly versatile biosensor specifically tailored for the spatio-temporal real time detection of proteolytic activity in plant systems, including Arabidopsis metacaspases. The modularity of the biosensor allows its implementation in various assays to identify and measure specific protease family activities through the inclusion of specific amino acids derived from a substrate cleavage sequence. Utilizing both in vitro and in vivo approaches, we demonstrate the biosensor’s efficacy in reliably detecting particular proteolytic activities within plant cellular environments. The versatility of this biosensor offers promising advancements for the plant research community, offering the ability to distinguish different protease activities in real time and providing enhanced spatial resolution of proteolytic events within plant systems.

## RESULTS

### FRET reporters can efficiently report proteolysis in plants

To engineer a Förster resonance energy transfer (FRET) sensor, incorporating a protease cleavage site between the two fluorophores, we selected mNeonGreen and mRuby3 as a well characterized and effective FRET donor-acceptor pair ^45^. This pair has a high quantum yield, large Stokes shift, fast maturation, high photostability, displaying improved sensitivity as a green/red fluorescent protein combination, and already successfully implemented in a biosensor for the detection of histidine kinase activity in bacteria and in plants to study Receptor-like kinase interactions ^45,46,47^Click or tap here to enter text..

To test the functionality of our FRET biosensor concept, we cloned a Tobacco Etch Virus (TEV) recognition site (TEVrs) in between the two fluorophores and named this sensor FRET^TEVrs^. TEVrs generally contains ENLYFQ↓G (↓: indicates the cleavage site) with a relative flexibility at the P1’ position and is cleaved by the highly substrate specific TEV NIa protease (TEVp). By engineering the TEVrs in between the two fluorophores they can be released from their proximal confirmation upon cleavage by TEVp (Fig. 1A). Our FRET^TEVrs^ sensor was recombinantly expressed and purified to high yield using bacteria as heterologous production system (Suppl. Fig. 1A). FRET^TEVrs^ underwent effective processing upon co-incubation with TEVp *in vitro* at equimolar levels (Fig. 1B). The processing of FRET^TEVrs^ *in vitro* became more pronounced at increasing concentrations of TEVp. (Suppl. Fig. 1B). Notably, the FRET ratio (mRuby3/mNeonGreen) displayed a significant reduction quantified through a plate reader, confirming the functionality of the biosensor (Fig. 1C). These findings validate the functionality of our biosensor and justify the selection of the mRuby3/mNeonGreen as FRET pair, emphasizing its suitability for subsequent experimental applications.

**Figure 1.**
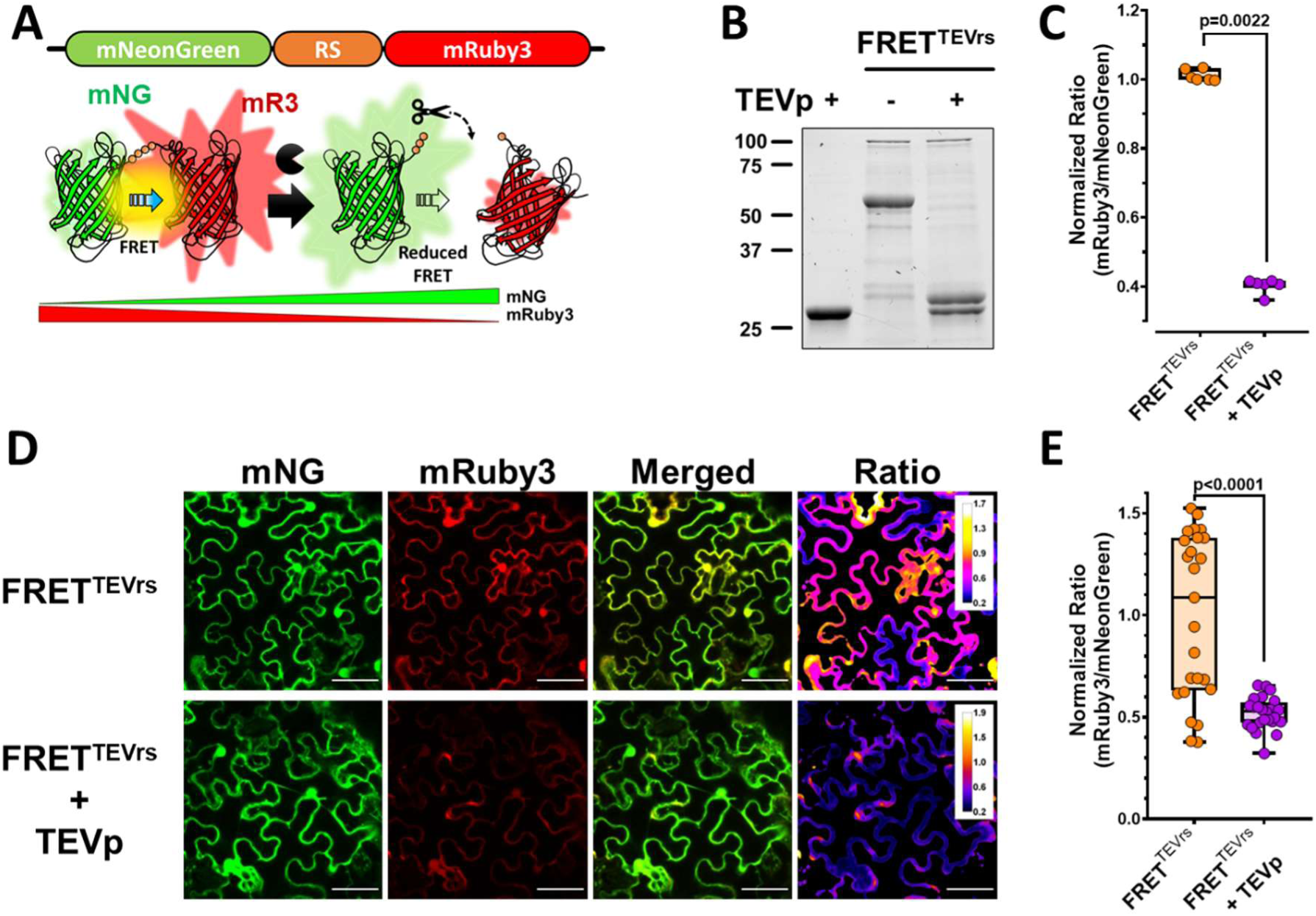
FRET sensor for the detection of protease activities in plants. **A)** Conceptual design of the FRET reporter for the detection of proteolytic activity. mNG: mNeonGreen; RS: recognition site and variation of relative donor/acceptor fluorescent levels. **B)** SDS-PAGE gel showing recombinant FRET^TEVrs^ purified sensor processing *in vitro* by incubation with TEVp. **C)** Fluorescent quantification of protease activity *in vitro* measured in a plate reader (p-values report nonparametric Mann–Whitney test, n=6 per condition). **D)** Panel displaying confocal images showing individual mNG and mRuby3 and merged fluorescent levels of FRET^TEVrs^ (*p35S:: FRET^TEVrs^ ::35St*) in presence absence of TEVp (*p35S::TEVp*) *N. benthamiana*. The last panel displays the fluorescent level ratio calculated in fire range (Bars represent 50 μm). **E)** Quantification of the normalized acceptor/donor level ratio (mRuby3/mNeonGreen) of FRET^TEVrs^ confocal images expressing the sensor alone (orange) and the sensor in combination with TEVp (purple). (p-values report two-tailed nonparametric Mann–Whitney test, n=25 per condition).

In the pursuit of enhancing FRET efficiency, we explored the influence of weak helpers, a strategy previously documented for intra- and intermolecular FRET ^48^. To this end, we subcloned our sensor and introduced weak helper peptides “WW” and “Wp1/2” at the N- and C-termini of FRET^TEVrs 48^. Our data revealed that the sensors were processed *in vitro* and reported a reduction of FRET emission when incubated with TEVp in an indistinguishable manner between the different FRET versions (Suppl. Fig 1D-E). Yet, we observed a statistically significant effect on FRET transmission in normal conditions with the WW-Wp1 pairs (Suppl. Fig. 1C), and we maintain this configuration with the WW and Wp1 helpers at the N- and C-terminal ends in subsequent biosensor design.

To assess the functionality of the reporter *in vivo*, the FRET^TEVrs^ construct, incorporating the optimized WW-Wp1 pair helpers, was transiently expressed under the control of the 35S promoter in *Nicotiana benthamiana*. This expression was carried out both in the absence and presence of TEVp, which was also expressed under the control of the same promoter. Upon co-expression with TEVp, a notable reduction in mRuby3 fluorescence was observed as shown by the decrease of the mRuby3/mNeonGreen ratio values (Fig. 1D). Quantification of the relative fluorescence values in cytosolic areas confirmed a reduction of the acceptor fluorescence to 50% in presence of TEVp in comparison to sample without protease (Fig. 1E). This decrease in FRET ratio provides robust evidence of the effective proteolytic processing of the FRET^TEVrs^ sensor *in vivo* by the TEVp, validating its functionality within a living plant system.

### FRET-based proteolysis reporters can detect viral infections in plants

We assessed whether the FRET^TEVrs^ sensor can work as a reporter during viral infection. The commonly used TEVp is an optimized, and in some cases engineered, protein from the Tobacco Etch Virus Nuclear-inclusion-a endopeptidase (NIa). TEV affects plant development and growth ^49^ and the protease in TEV NIa is required for cleavage and functionality of the original viral polyprotein chain into seven functional proteins^50^. To test this, we used 6–8 weeks-old plants of *N. tabacum* inoculated with TEV or healthy plants prior visible signs of viral infection (Suppl. Fig. 2). The FRET^TEVrs^ sensor was infiltrated in leaves of untreated and virus inoculated plants displaying slight differences in the fluorescent values (Fig. 2A). The FRET ratio records in infected plants were reduced and more heterogeneous than in healthy plants, with patches of adjacent cells displaying highly reduced values. Quantification of the FRET showed significant differences between the healthy and virus-inoculated plants (Fig. 2B). The quantification trends in infection plants indicates that there are likely two populations of data, one showing similar levels to healthy plants and a subset of cells with reduced FRET rations, with similar values to plants overexpressing TEVp. This phenomenon could be explained by the heterogeneity of the virus infection across different tissues of the plant, which results in varying levels of viral presence and activity. Alternatively, it may be due to the decreased activity of the endogenous viral NIa protease compared to the overexpression of TEVp achieved through transient transformation. These results further highlight the robustness of our engineered FRET biosensor, demonstrating its capability to respond to proteolytic activities within a biological context.

**Figure 2:**
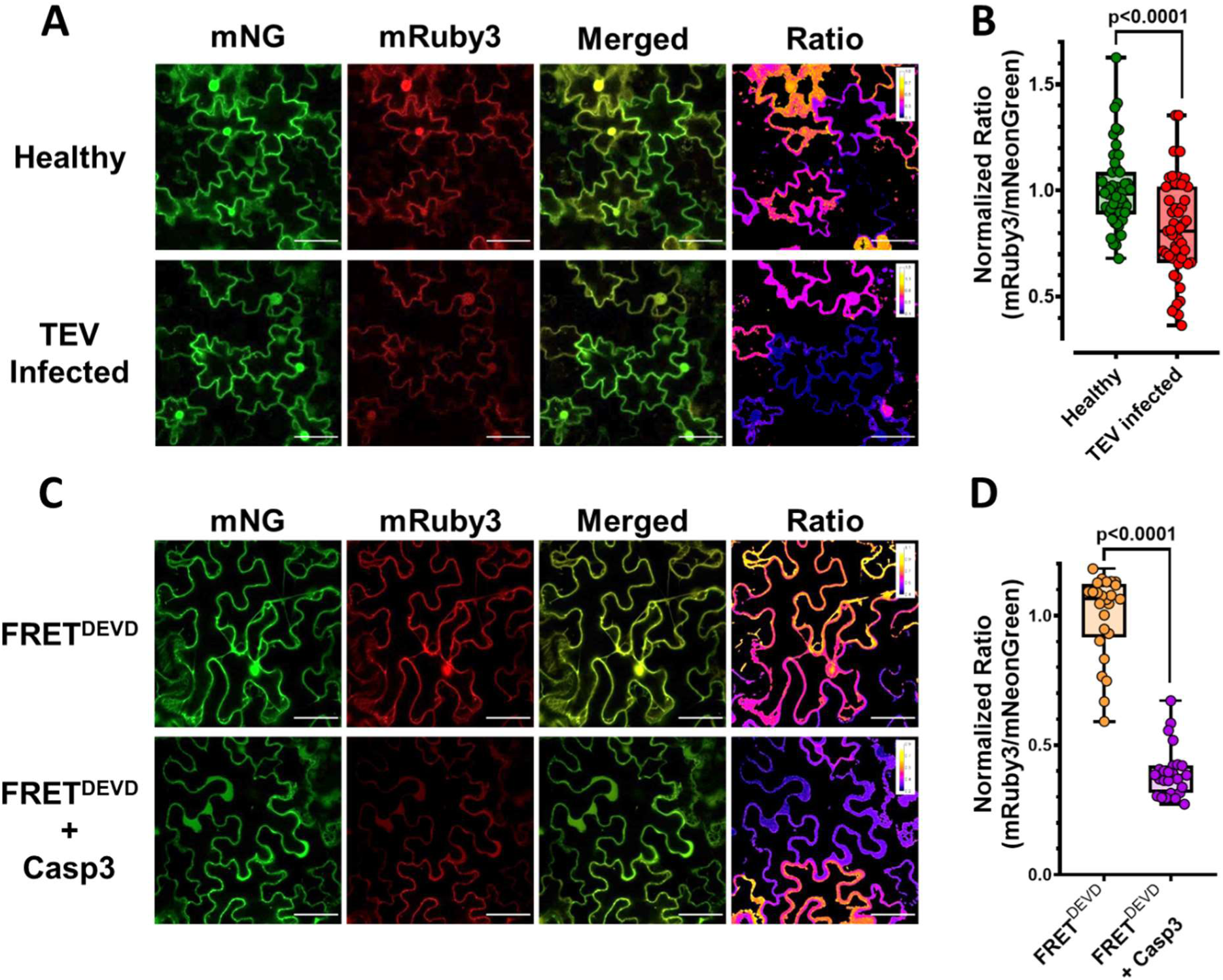
FRET biosensors can be used for the detection of different proteolytic activities. **A)** FRET^TEVrs^ can detect TEV presence in *N. tabacum* by reporting its protease activity. Panels show confocal images for the fluorescent of mNeonGreen, mRuby3, merged channels and ratio (Bars represent 50 μm). **B)** Normalized quantification of the ratio signal acceptor/donor (mRuby3 FRET/mNeonGreen emission) of FRET^TEVrs^ for images obtained in non-infected plants (green) and infected with TEV (red) showing significant differences between samples (p-values report two-tailed nonparametric Mann–Whitney test, n=50 per condition). **C)** Confocal images of transient expression of FRET^DEVD^ sensor in presence or absence of caspase-3. Separated channels for mNeonGreen, mRuby3, merged and ratio picture (Bars represent 50 μm). **D)** Normalized quantification of the ratio signal acceptor/donor (mRuby3 /mNeonGreen emission) of FRET^DEVD^ in leaf areas with presence and absence of transiently expressed caspase-3 (orange and purple respectively; p-values report two-tailed nonparametric Mann–Whitney test; n=25 per sample).

### FRET-based protease reporters can be engineered for the detection of different proteolytic activities

To test the sensor versatility, we sub-cloned the active domain of mammalian caspase-3 ^51^ into a plant compatible expression system. Caspase-3 is a widely studied protease and there exists multiple tools to test its activity, including chemical fluorescent probes. An example is CR(DEVD)2, a fluorescent dye that increases fluorescence by proteolysis upon cleavage by caspase-3. Also, CR(DEVD)2 has been previously used in plants to detect caspase-3-like activity in pollen and vegetative tissues, showing its cellular permeability ^52,53^. Upon transient expression of caspase-3 in leaves of *N. benthamiana,* CR(DEVD)2 successfully reported high fluorescent levels indicating that the transiently expressed caspase-3 is active in exogenous systems like plants (Suppl. Fig. 3A-B). This result encouraged us to test caspase-3 as additional protease to adapt our FRET sensors. We cloned the caspase-3 minimal substrate sequence DEVD in the protease recognition site among WW-mNeonGreen and mRuby3-Wp1 ^54^ and named the sensor FRET^DEVD^. Confocal imaging showed a reduction of the mRuby3 FRET signal when caspase-3 was co-expressed with the sensor in *N. benthamiana* leaves (Fig. 2C). Quantification of the fluorescence levels in planta revealed a significant decrease in the FRET ratio when caspase-3 was present (Fig. 2D). The FRET values of the sensor in presence of the protease showed a reduction to 40% in relation to the total values without protease. The data obtained is very similar to the FRET reduction obtained with FRET^TEVrs^ and TEVp overexpression above, and they fell within the same range of other studies testing caspase-3 activity in mammalian cells^48^. Capase-3 and caspase-3-like activities have been linked in plants to cell death events ^12,55^, and some plant proteases are accounted to contribute to cell-death symptoms. To avoid possible side effects of cell death, we imaged at early timepoints after infiltration (2 days after infiltration) as we could observed clear signs of cell death in leaves from day 6, likely due unspecific cleavage of highly expressed caspase-3 in *N. benthamiana* (Supp. Fig.3C).

### Adaption of the FRET sensor towards specific metacaspase activity

Next, we extended our investigation by exploring modifications to the protease recognition site (RS), enhancing the sensor adaptability and the potential applications in studying proteolytic processes in plant systems. Among plant proteases, metacaspases are one of the most extensively studied groups. Metacaspases play crucial roles in various physiological processes, including programmed cell death, stress responses, and developmental pathways^18,19,22,23,28,56–58^. In addition, plenty of information exist on metacaspases activity regulation, inhibition and a handful of *bona fide* substrates have been identified to date ^19,26,59–61^. Previous studies showed the preference of metacaspases for processing after arginine and lysine amino acids ^19,29,61,62^. This information allowed us to tailor our sensor by inclusion of metacaspases substrate sequences in the recognition site and test their function upon exposure to active metacaspases, including physiological relevant conditions.

We incorporated metacaspases minimal recognition motifs VRPR and PROPEP1-derived sequences VTSRATKV and VTSRATKVKAKQRG named PROPEP1 CLEAVAGE SITE 1 and 2 (PCS1 and PCS2 respectively) obtaining FRET^VRPR^, FRET^PCS1^ and FRET^PCS2^ sensors (Fig.3A). Structural predictions of the FRET^VRPR^ sensor showed that the recognition site is highly exposed while the fluorescent proteins maintain proximity (Suppl. Fig. 4A).

**Figure 3.**
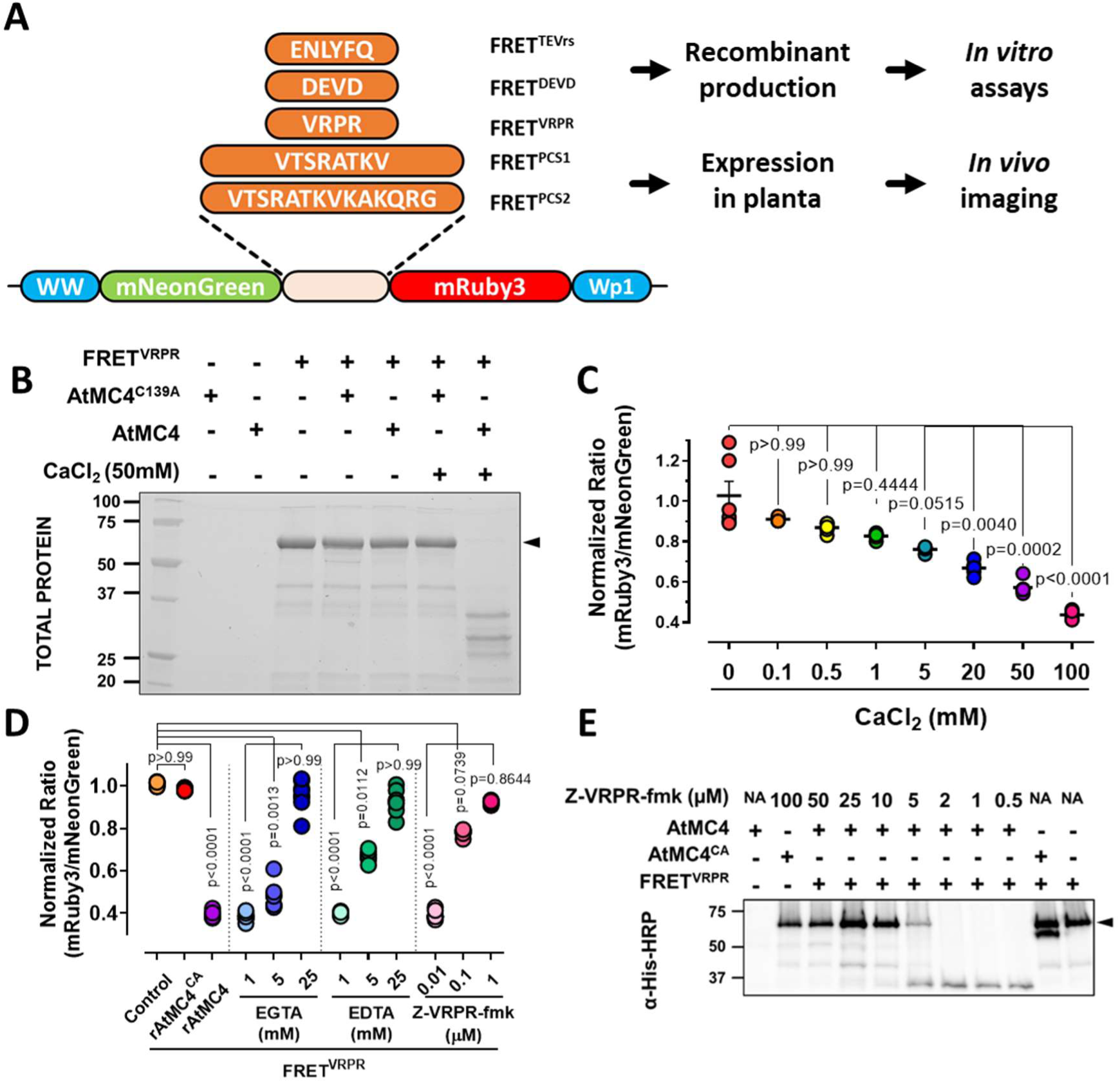
Characterization of metacaspase specific FRET sensors. **A)** Schematic design of different FRET sensors. The protease recognition site (RS) show the permutation of amino acids as minimal substrates to confer specificity to different proteases, including FRET^TEVrs^, FRET^DEVD^, FRET^VRPR^ and PROPEP1-based substrate sequences FRET^PCS1^ and FRET^PCS2^. In blue the weak helpers WW and Wp1 are indicated. **B)** Total protein gel depicting calcium requirement for AtMC4 processing of FRET^VRPR^ in vitro. Arrows indicate full-size FRET^VRPR^. **C)** Co-incubation of AtMC4 and FRET sensor at increased levels of CaCl2 reveal a dose dependent dosage for the activation of AtMC4 in the mM range by measuring ration values of mRuby3/mNeonGreen of FRET^VRPR^ (p-values report Kruskal-Wallis test results, n=8). **D)** Protease drop of FRET^VRPR^ ratio values are effectively recovered upon application of known inhibitors of AtMC4 *in vitro*. Increasing levels of EDTA/EGTA and Z-VRPR-fmk results in reconstitution of fluorescent, ultimately reaching levels similar to unprocessed sensor (p-values report Kruskal-Wallis test results, n=8). **E)** Western Blot showing abolition of FRET^VRPR^ cleavage, evidencing a dose dependent effect of the inhibitor Z-VRPR-fmk.

Recombinantly expressed FRET^VRPR^ displayed processing *in vitro* when incubated with active AtMC4 in presence of calcium (Fig. 3B, Suppl. Fig. 4A-B). Moreover, the FRET values were reduced in a dose-dependent manner upon presence of increasing values of calcium (Fig. 3C).

Upon testing with other type-II metacaspases, FRET^VRPR^ reported similar changes in its FRET ratio value than to AtMC4 (Suppl. Fig. 4D-G). Next, we tested whether FRET^VRPR^ processing by AtMC4 could be reverted by application of known metacaspase inhibitors. FRET loss caused by proteolytic cleavage was reduced in presence of calcium chelators EDTA and EGTA by blockage of AtMC4 self-activation. At 25 millimolar levels of these chelators, we could not distinguish an effect of AtMC4 on the recorded values for FRET^VRPR^ in comparison to FRET^VRPR^ alone. Moreover, when a metacaspase-specific inhibitor (Z-VRPR-fmk), was included in the reaction, the ratio values were restored to normal levels at micromolar concentration of the inhibitor (Fig. 3D). *In vitro* test of the sensor performance using Z-VRPR-fmk abolished FRET^VRPR^ degradation products at similar concentrations that FRET ratio was restored (Fig3E). Our findings indicate that FRET^VRPR^ sensor undergoes processing by metacaspases in a calcium dependent manner and that the reporter functions can be abolished by using metacaspase specific inhibitors. Importantly, we also reported that fluorescent measurements reliably reflect protease processing. AtMC9, a calcium-independent protease, demonstrated the ability to process the FRET^VRPR^ sensor, as evident from the reduction in mRuby3 values, establishing plate reader values as the reference standard. Additional sensors based on PROPEP1 processed sequences, (FRET^PCS1^ and FRET^PCS2^) responded similarly to FRET^VRPR^ upon testing their function in presence of AtMC4 (Suppl. Fig. 5A-B).

### Metacaspase FRET sensors report proteolytic activity *in vivo*

We further investigated the functionality of our reporter by its expression in plants, for the detection of plant proteases action *in situ*. Previous studies have shown that calcium released from wounded tissues can activate AtMC4 in nearby cells ^19^. Thus, we tested whether our FRET reporters can serve as a tool reporting metacaspase proteolysis in real time after damage. We tested different FRET sensors in the presence of AtMC4 in *N. benthamiana* (Suppl. Fig. 6). In plants transiently co-expressing AtMC4 and FRET^PCS1^ in *N. benthamiana* leaves, a noticeable reduction in FRET signal levels became evident in cells adjacent to damaged areas. (Fig. 4A). The continuous decrease of the signal after laser ablation and in proximity denotes that the effect is not due to photobleaching but rather a FRET changes from the sensor over a period of five minutes (Fig. 4B; Suppl. Video 1). Moreover, the time range at which the FRET reporter decreases aligns with the expected dynamics at which AtMC4 processes PROPEP1 in *Arabidopsis* roots. Contrarily, the FRET ratio of a reporter containing a linker of Glycine and Serine at the recognition site (FRET^6xGS^) did not show any visible FRET change (Fig. 4B, Suppl. Video 2). Consistent with the earlier findings, these results suggest that FRET reporters, in this case FRET^PCS1^, can be designed for the detection of protease-specific activities in their natural environment. This customization can be achieved by exchanging the sequences within the recognition sites for specific substrates to a certain protease or avoiding their processing by inserting non-specific sequences.

**Figure 4:**
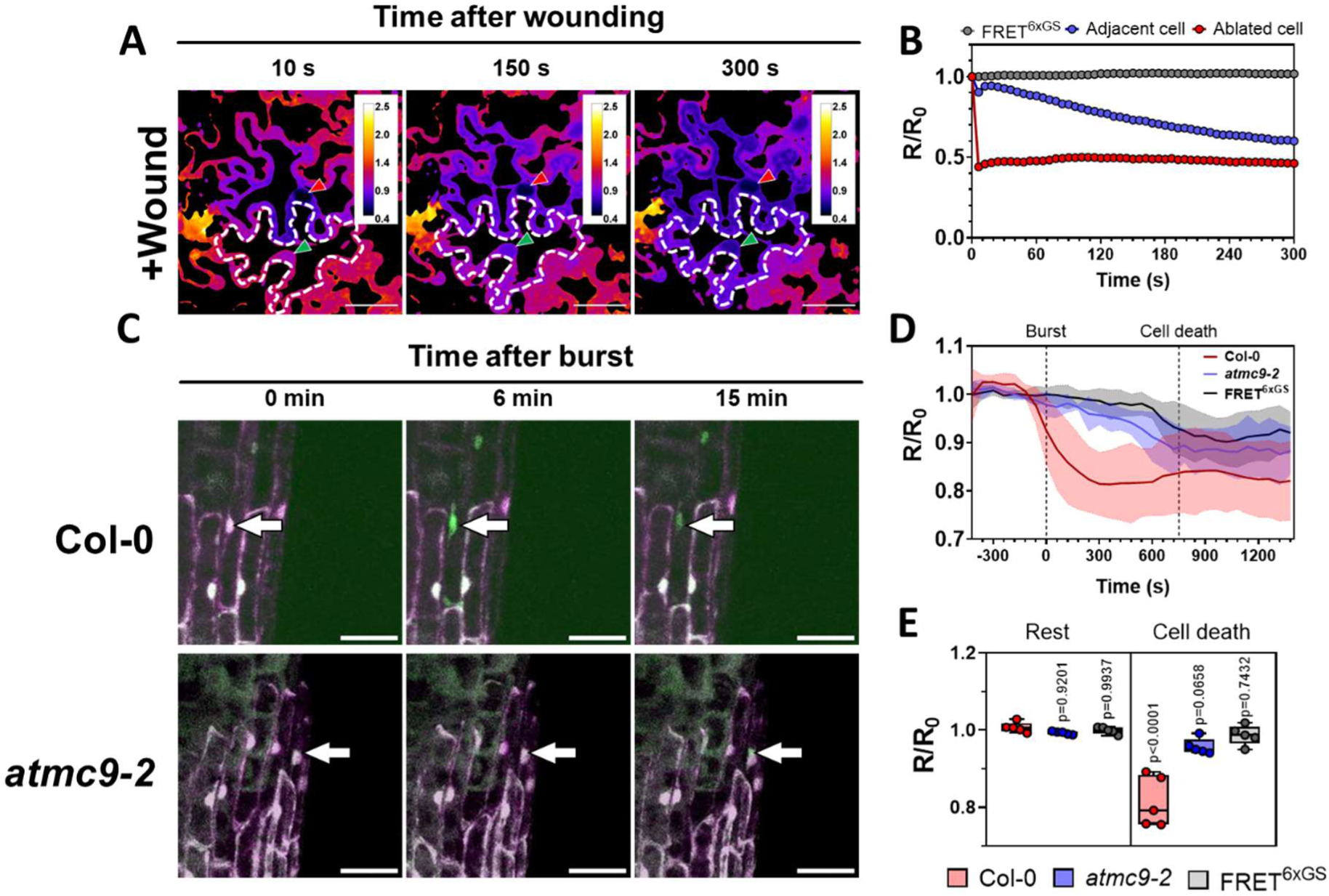
Metacaspase-tailored FRET sensors report protease activity of the proteases in biological contexts. **A)** FRET ratio of *N. benthamiana* cells after wounding transiently co-expressing FRET^PCS1^ and AtMC4 (*p35S::AtMC4*). Images were taken for a time-lapse and ratio at timepoints at 10,150 and 300 seconds are displayed. Cell marked with red arrow was damaged using laser ablation, resulting in immediate drop of fluorescence. The adjacent cell, indicated with the dotted line and the green arrow show a gradient and consistent drop in the ratio levels after wounding. **B)** Time-lapse quantification of the FRET^PCS1^ ratio of cells after damage. Values plotted corresponding to damaged cell by ablation (red), adjacent cell (green), and ratio values of an adjacent cell to damage using a negative control with a GS linker FRET^6xGS^ (blue). **C)** FRET^VRPR^ expression in Arabidopsis root cap cells. Arrows indicate the last layer of root cap prior to cellular collapse in Col-0 and *mc9-2 mutant lines.* **D)** Plot showing the normalized values of Arabidopsis plants expressing FRET^VRPR^ signal during cellular burst and cellular collapse in Col-0 and *mc9-2* backgrounds and FRET^6xGS^. **E)** Quantification of FRET^VRPR^ values of cells undergoing cell death at resting timepoints (-120 s before burst) and 300 s after cells were detected to enter bursting process (p-values report two-tailed nonparametric Mann–Whitney test in relation to the normalized ratio of Col-0 in resting conditions, n=5).

*Arabidopsis* plants transformed with FRET^VRPR^ or FRET^6xGS^ were tested for the detection of metacaspase proteolytic activity. In previous studies, AtMC9 was reported to be expressed in the root cap, xylem, and lateral root poles with direct association with programmed cell death events ^17,61,63^. We centered on the developmental process within the root cap and monitored the fluorescent levels of the FRET sensors in both wild-type and AtMC9 mutant plants (*atmc9-2*). FRET values showed a reduction in mRuby3 fluorescent values in specific cells undergoing the final stages of programmed cell death, prior cellular collapse (Fig 4. C). Our results hints towards a distinguishable activation of metacaspase activity occurring at late stages preceding developmental programed cell death of lateral root cap. Quantification of the cells undergoing this process presented detectable FRET changes in wild type plants (Fig. 4D). Interestingly these values were significantly reduced in *mc9-2* mutant lines, demonstrating the essential role of AtMC9 to execute metacaspase activity in root caps, as indicated by our reporter FRET^VRPR^. Root cap cells experienced a rapid collapse within a short timeframe of minutes. This collapse was accompanied by the simultaneous loss of fluorescence signals from both the donor and acceptor of the FRET reporter, signifying the complete cessation of cellular activity. (Suppl fig. 7). Our findings are consistent with previous reports on developmental programmed cell death in Arabidopsis, where cell permeabilization, a marked increase in hydrolase enzymatic activity, and subsequent vacuolar collapse collectively result in the efficient clearance of cellular debris. ^64^. However, the precise physiological role of AtMC9 in developmental programmed cell death within the root cap is yet to be comprehensively examined. Our reporter offers a platform for hypothesis, proposing that signals of presently unknown nature and origin might precede cell death, proving sufficient to activate AtMC9 at this specific instance and location.

## DISCUSSION

Here we describe a tailor-made FRET reporter for the detection of protease activity, focusing on proteolytic events occurring in planta. We show that the sensor can report specific proteolytic activities in real-time and with spatial resolution in the presence of active proteases. Moreover, we demonstrate that the FRET sensor values are conditioned to the intrinsic properties of the protease tested, as it was shown for the calcium activated AtMC4 *in vitro* and *in vivo* experiments during wounding events. Our results indicate that the FRET ratio levels directly correlate to specific proteolytic activity and can report levels of activity by the proteolytic machinery *in vivo*.

The findings revealed in this study suggest that mRuby3 and mNeonGreen work efficiently as acceptor and donor pair in intramolecular FRET systems in plants. This evidence confirms previous reports where the same fluorescent proteins have been used ^45,65^. Furthermore, the detection of protease activity led to a reduction of 40-50% in comparison to resting-state values, a magnitude akin to that observed with FRET sensors reporting proteolysis in other systems ^48^. The photostability of both proteins allows for detection of long timelapses at different timepoints during events like wounding in transient expression or *Arabidopsis* root timelapses. While this FRET pair suffices for our current purposes, other studies may benefit from utilizing alternative FRET pairs with improved transmission or fluorescent emission signals that exhibit less overlap with endogenous autofluorescent levels. We enhanced the system by incorporating weak helpers, which led to an increase in basal fluorescence levels. Surprisingly, the sensors reported cleavage in a remarkably similar manner to a FRET sensor without helpers. This suggests potential advantages in situations where obtaining sensor expression is challenging, while still preserving its ability to accurately report peptidase activity.

Our initial results using FRET^TEVrs^ and overexpression of TEVp strongly support the capacity of this reporter to detect other proteases with biological relevance in plants. In fact, when *N. tabacum* plants were infected with TEV, the sensor was capable of efficiently reporting proteolytic activity from NIa protease from viral inoculated plants. Numerous viruses encode proteases in their genome with the function of processing polyproteins into functional, active proteins necessary for infection. For example, the Nuclear Inclusion a proteases from Turnip Mosaic Virus or Potato Yellow Virus cleave after VXXE/VXXQ where X has a flexibility for different amino acids.^66,67^. Being viruses are one of the biggest causes of crop losses worldwide ^68^, specific proteolytic sensors could be used to detect viral presence before plants show evident symptoms of infection. Therefore, protease sensors could be another invaluable resource to identify viral presence or being used as screening readout at identifying inhibitors that effectively block the proteolytic activity of a given viral protease. This application could be translated to other systems like blockage of viral protease affecting humans with known substrate affinities as done for SARS-CoV-2 main protease ^69^.

Through sensor customization, we extended our detection capabilities to include caspase-3 activity in plants. By swapping the TEV recognition site to DEVD and testing, we efficiently reported caspase-3 activity in plant systems. Multiple studies in plants referred to caspase-3-like activity and DEVDase activity. These studies employed tools from other fields to test caspase activities in plants without specific protease assignments, indirectly linking them to cell death. Our results indicate that caspase-3 activity in plants can be measured with our FRET^DEVD^ sensor. However, we refrain from speculating that caspase-3 induced cell death in plants occurs through a conserved mechanism with mammals, given the absence of real caspase orthologs in plants. We suggest that the observed cell death may result from prolonged overexpression of a highly active protease. This overexpression could result in the non-specific cleavage of essential cellular components, leading to the malfunction of crucial cellular processes and ultimately culminating in cell death.

We also used the sensor to create metacaspase specific activity reporters. The sensor effectively reported different metacaspases processing *in vitro*. Moreover, we could prevent the fluorescent changes by using known inhibitors of metacaspase activity. Different calcium chelators were reported to hamper the initial self-processing of AtMC4 functions *in vitro* and *in vivo* ^19,26,29,30,62,70^. Similarly, the sensor showed high responsiveness to EGTA and EDTA inhibitors blocking AtMC4 action. Z-VRPR-fmk specific metacaspase inhibitor contains a VRPR sequence mimicking metacaspase substrates and a warhead that covalently binds to the active site, effectively inhibiting its hydrolytic activity, rather than blocking its self-processing. Its application Z-VRPR-fmk at micromolar concentrations managed to reduce metacaspase activity by decreased mRuby3/mNeonGreen ratio and abolition of its cleavage as seen by immunoblots. Taken together, our *in vitro* results suggest that the FRET^VRPR^ performs effectively under conditions known to activate metacaspases, with the ability to be reversed by the application of inhibitors.

These results were confirmed *in vivo* by addressing metacaspase activity after wounding. Cell ablation sufficed to induce a systemic transient calcium wave and a local centralized accumulation of calcium. Our results upon damage showed FRET ratio reduction in adjacent cells to the targeted area in a timeframe of 5 minutes after wounding. Our results go back-to-back with the expected timeframes at which PROPEP1 is released from the tonoplast in *Arabidopsis* roots upon laser-ablation treatments. Moreover, root imaging in stably expressed lines indicates that there is a certain mechanism that activates AtMC9 in the latter events of programmed cell death. While the importance of this activity remains unclear, specific proteolytic reporters like FRET^VRPR^ might serve to characterize the time and events at which some proteases are active.

In summary, we successfully developed a series of advanced proteolytic reporters capable of detecting various proteolytic activities. Genetically encoding reporters offers the advantage of producing them recombinantly for biochemical characterization and transformation into model organisms. As we demonstrated, our FRET sensors are adaptable to specific protease requirements, allowing the customization of recognition sites according to the substrate of interest. Moreover, the versatility of genetically encoded sensors enables their targeting to specific sub-cellular compartments through the incorporation of targeting signals, making them ideal for situations where a particular protease is exclusively located in a specific organelle. FRET reporters exhibit a notable speed advantage over other biosensors, providing an immediate readout of activity, while some biosensors require maturation before fluorescent or luminescent signal changes. Additionally, FRET reporters offer the advantage of containing two layers of fluorescence information, facilitating normalization even under conditions where expression may be suboptimal. Lastly, genetically encoded biosensors play a crucial role in delineating proteolytic dynamics over time and space for a given protease, allowing us to pinpoint the specific window of time during which it becomes active and explore conditions that may trigger its activity.

## MATERIAL AND METHODS

### Plant material and growth conditions

7 to 10 days-old *Arabidopsis thaliana* (Col-0) plants were used for root imaging. Plants were sterilized using the gas chlorine method and placed in squared plates containing ½ MS solid media, covered with aluminum foil and stratified during 3-4 days at 4 °C in the darkness. After stratification, plants were placed for 10 days in a growing chamber at 21 °C with 16-8 hours of light/dark cycle at a light photo-intensity of 100 μmol · m^-2^· s^-^1. Four to six weeks old *Nicotiana benthamiana* (Nb1) plants were used for transient expressions. TEV inoculated plants were ordered at the Deutsche Sammlung von Mikroorganismen und Zellkulturen (DSMZ) Institute. Freeze-dried plant material was placed in a precooled mortar and a few drops of inoculation buffer was added (50 mM sodium/potassium phosphate buffer adjusted to pH 7.0, 1 mM EDTA, 5 mM diethyldithiocarbamate, 5mM thioglycolic acid) and ground to a thick paste. Inoculation buffer is added to a final volume of 4 ml and rubbed in a maximum number of three plants. *Nicotiana tabacum* were used as host plants for the TEV inoculation, grown at 22-25°C in the greenhouse at 14-10 hours of light/dark cycle. Healthy and infected plants were grown in the same chamber.

### Cloning and transformation

#### Cloning of FRET^TEVRS^ concrete His tag?

We used GreenGate collection constructs (Lampropoulos et al., 2013). TEVp original sequences were ordered from addgene (pcDNA3.1 TEV: Plasmid #64276). The DNA fragment corresponding to WW sequence was custom synthesized (Integrated DNA Technologies). Original caspase-3 plasmid (Casp3p30) was obtained from Prof. Savvas Savvides containing the p12 and p18 caspase 3 subunits rendering a total p30 size protein that is constitutively activated. To generate GreenGate compatible entry clones we used collection compatible entry plasmids or CloneJET PCR Cloning Kit (Thermo Scientific) according to the manufacturer manual. Primer names and sequences used in this study can be found in Suppl. Table 1. Final plasmid construction by restriction and ligation was performed according to Lampropoulos *et al.*, (2103) using periods of 2 minutes for the ligation at 16 °C followed by enzymatic restriction during 2 minutes at 37°C for a total of 25 cycles. Ligation product was directly used for *E. coli* transformation and growing colonies were verified by colony PCR and Sanger sequencing. All the plasmid details and names are indicated in Suppl. Table 2. Plasmids were transformed in *Agrobacterium tumefaciens* C58C1 and selected using the appropriate resistances.

### *N. benthamiana* transient transformation

*Agrobacterium tumefaciens* strains were grown in flasks containing 10-20 ml YEB media with their corresponding resistances (Gentamicin, Rifampicin and Spectinomycin) at 28 °C for 2-3 days with shaking. Bacterial concentration was estimated and the culture volume for a final OD600=1.5 spined-down and resuspended in the infiltration buffer (10 mM MgCl2, 10 mM MES buffer, 10 µM freshly added acetosyringone). Resuspended cells were incubated at 28 °C for 2-4 hours and infiltrated in 4 weeks old leaves of *Nicotiana benthamiana* or *Nicotiana tabacum*. Confocal imaging and tissue sampling was done at day 2 or 3 after infiltration unless specified on a Zeiss 710 confocal setup.

### Photo images

Photo images were taken in an adjustable custom-made stand with light bars using a Nikon Eos 650D photo camera.

### Confocal imaging

Seedlings with good expression levels were selected to T3 generation and imaged using a Zeiss LSM 710 confocal microscope. Plants were imaged using a 488 nm laser and adjusting the filters for GFP emission and PI staining. In our hands, the Zeiss 488 nm laser induced less noise ratio when imaging mNeonGreen, than the 512 nm laser and therefore we used this laser for FRET detection. Detection of the signal and gain for each channel were maintained constant to compare images. Gain changes in one channel were corresponded for the equivalent change in the next channel. Images were analyzed and edited using ImageJ free software. FRET ratio images were done as indicated for the donor and acceptor channels in Hander *et al.*, (2019).

Arabidopsis root images were acquired on an AxioObserver.Z1 inverted microscope coupled to an LSM900 point scanner (ZEN ver. 3.3.89.0008, Zeiss, Germany). Imaging was done with the Plan-Apochromat 20x/0.8 M27 objective, excitation was set at 488 nm and the two emission windows were 490 - 530 nm for mNeonGreen and 600-700 nm for mRuby3. For time lapse experiments, TipTracker was used for automatically tracking the growing root ^71^, images were collected every three minutes for up to 12 hours. Prior to analysis images (8-bit) were pre-processed in Fiji.

### DEVDase visualization with CR(DEVD)2

For the validation of the caspase activity, we used CR(DEVD)2 CV-Caspase 3 & 7 detection kit (Enzo Life Sciences, BML-AK118-0001). CR(DEVD)2 powder was dissolved in 100 µl dimethyl sulfoxide and diluted 1:5 in milliQ H2O prior to making a staining solution. The working solution was made by diluting 1:50 in milliQ H2O. Samples were incubated 30 minutes at room temperature before imaging. For confocal imaging, an excitation filter of 550 nm (540-560 nm) and a long pass >610 nm emission/barrier filter were used.

### SDS-PAGE and Immunoblotting

Protein samples were loaded in a stain-free 12,5% SDS-PAGE precast gels (BIORAD) in Tris-Glycine-SDS (TGS) buffer at a voltage of 200 V for 45 minutes approximately. Stain-free gels were activated and total protein on gel were detected and recorded using a ChemiDoc™ Imaging System (BioRad).

Gels were transferred to a precast TransBlot Turbo membrane for 3 minutes at 2.5 amperes and voltages up to 25 V in a BioRad membrane transfer system. Membranes were blocked in 5% skim milk in PBS-T (0,2% Tween) for 2 hours. The blotting of His-tagged proteins was done by 2 hours incubation in 1% skim milk with single step antibody conjugated to HRP (1:1,000) followed three washing steps for 10 minutes in PBS-T. Detection was performed by exposition to a (1:1) solution of Western Lightning® Plus-ECL reagents (Perkin Elmer) for 1 minute and chemiluminescence imaging in a ChemiDoc™ Imaging System (BioRad).

### Statistical Analysis

Statistical analyses were performed using GraphPad Prism 9.0 for Windows. As mentioned in the figure legends, statistical significances were assessed using nonparametric Kruskal–Wallis bilateral tests combined with post-hoc Dunn’s multiple pairwise comparisons; two-way nonparametric Student’s t-test Mann–Whitney test or ordinary one-way ANOVA and significant and non-significant p-values were indicated in the figures.

## Acknowledgements

We thank Prof. Dr. Savvas Savvides for sharing with us the plasmid containing the His-Casp3p30 sequence. The original *Nicotiana benthamiana* Nb1 seed stock was a gift from Prof. Gregory B. Martin. Funding: Wallenberg Academy Fellow 2021, <u>grant 2021.0071</u>; European Research Council 2021 Consolidator, grant 101044878 DETOXPEST; FWO14/PDO/166 to S.S.; the Ghent University Special Research Fund (grant 01J11311 to F.V.B.; Fund for Scientific Research-Flanders (FWO) grant (G021119N) and a Ghent University Concerted actions (GOA) grant (BOF19-GOA-004) to J.S; Fonds Wetenschappelijk Onderzoek – Vlaanderen (FWO, project grant G002620N), European Research Council (ERC CoG EXECUT.ER 864952) to M.K.N. and M.V.D.

---

**Supplementary Table 1.**
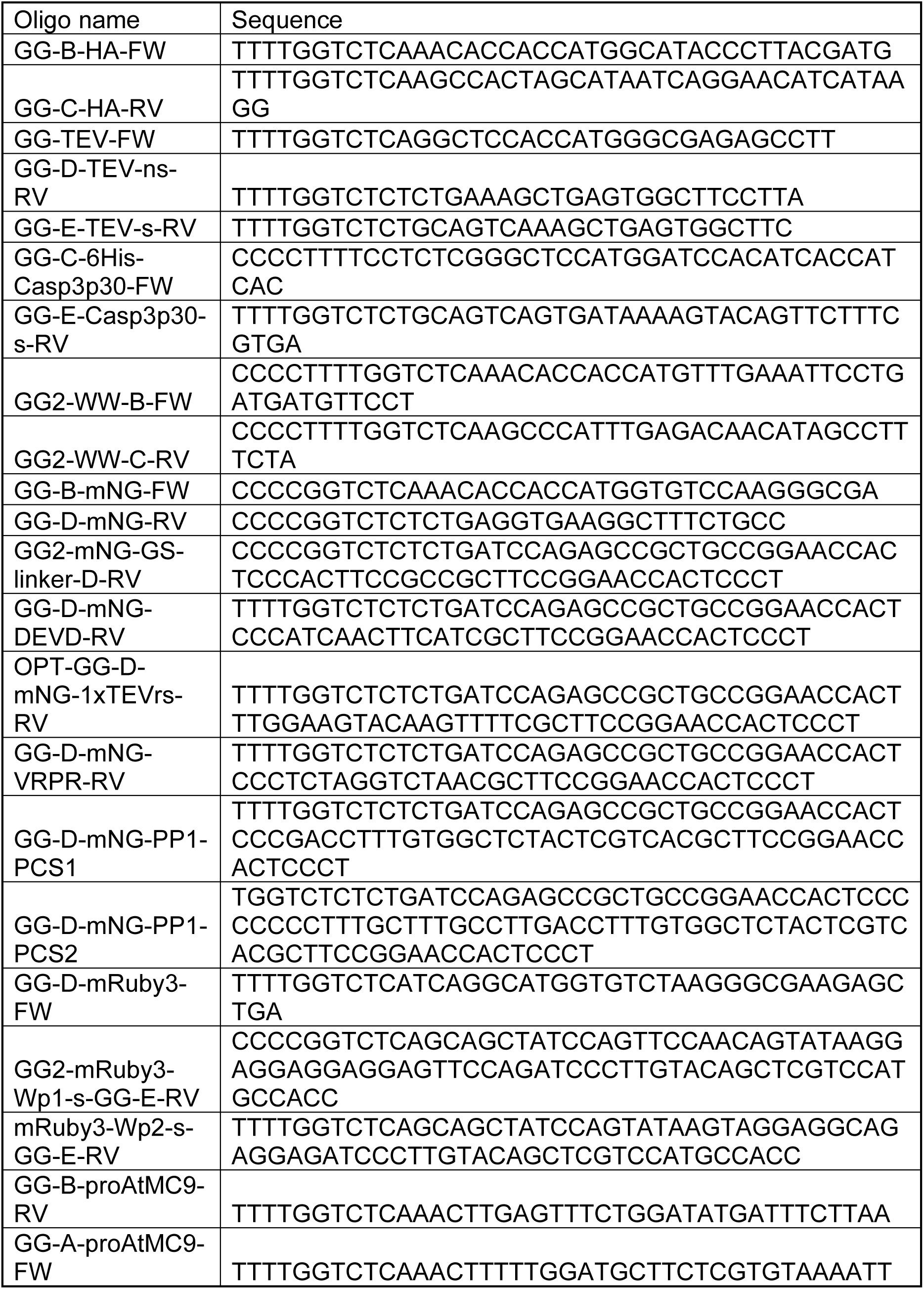
Oligo names and sequences used in this study.

**Supplementary table 2.**
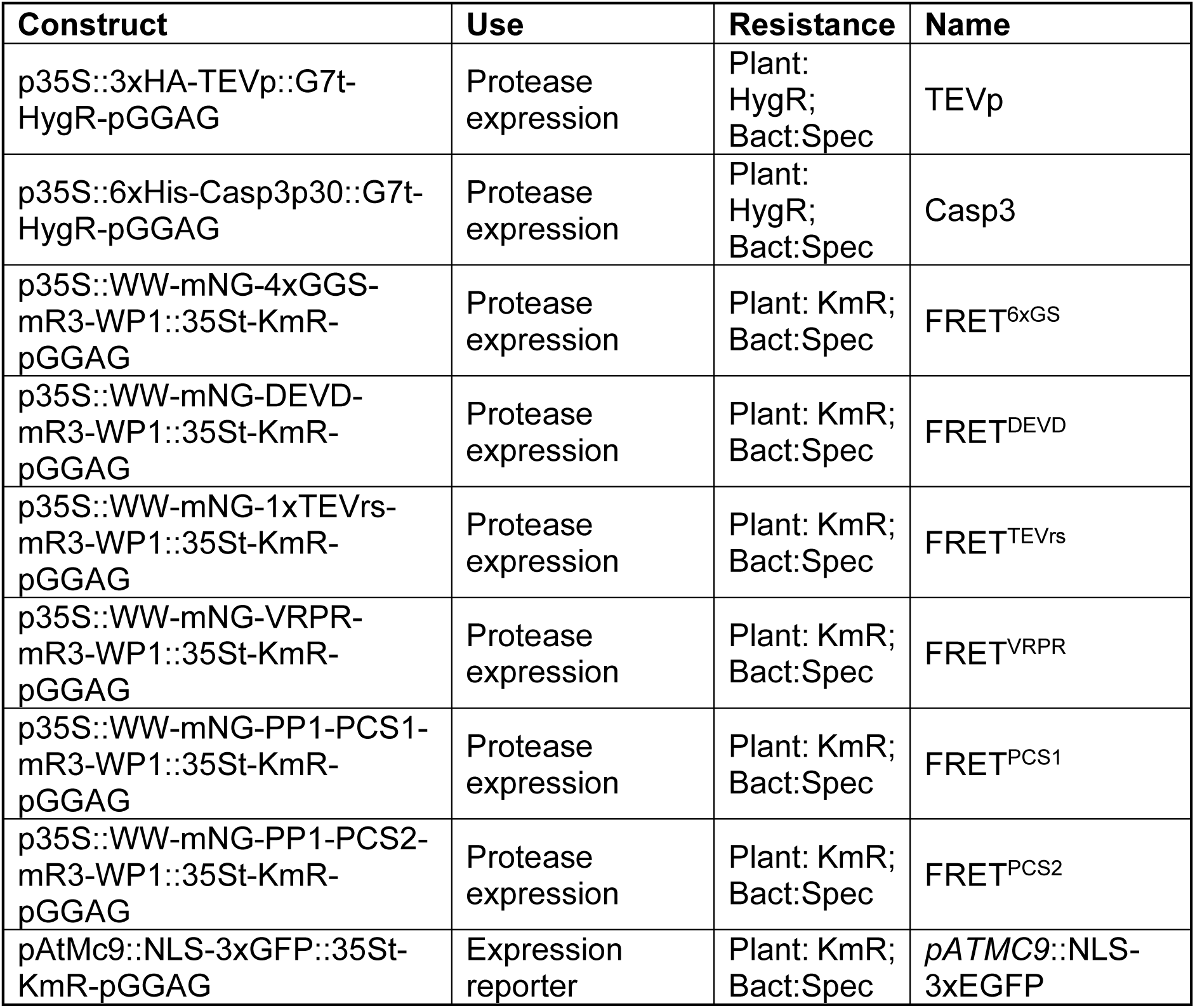
Constructs used in this study. Entry modules were obtained from the GreenGate collection or cloned (See Suppl. table 1). HygR: indicates plant resistance to Hygromycin; BarR: indicates plant resistance to BASTA; SdR: indicates plant resistance to Sulfadiazine; KmR: indicates plant resistance to Kanamycin; Spec: indicates bacterial resistance to Spectinomycin.

**Supplementary Figure 1.**
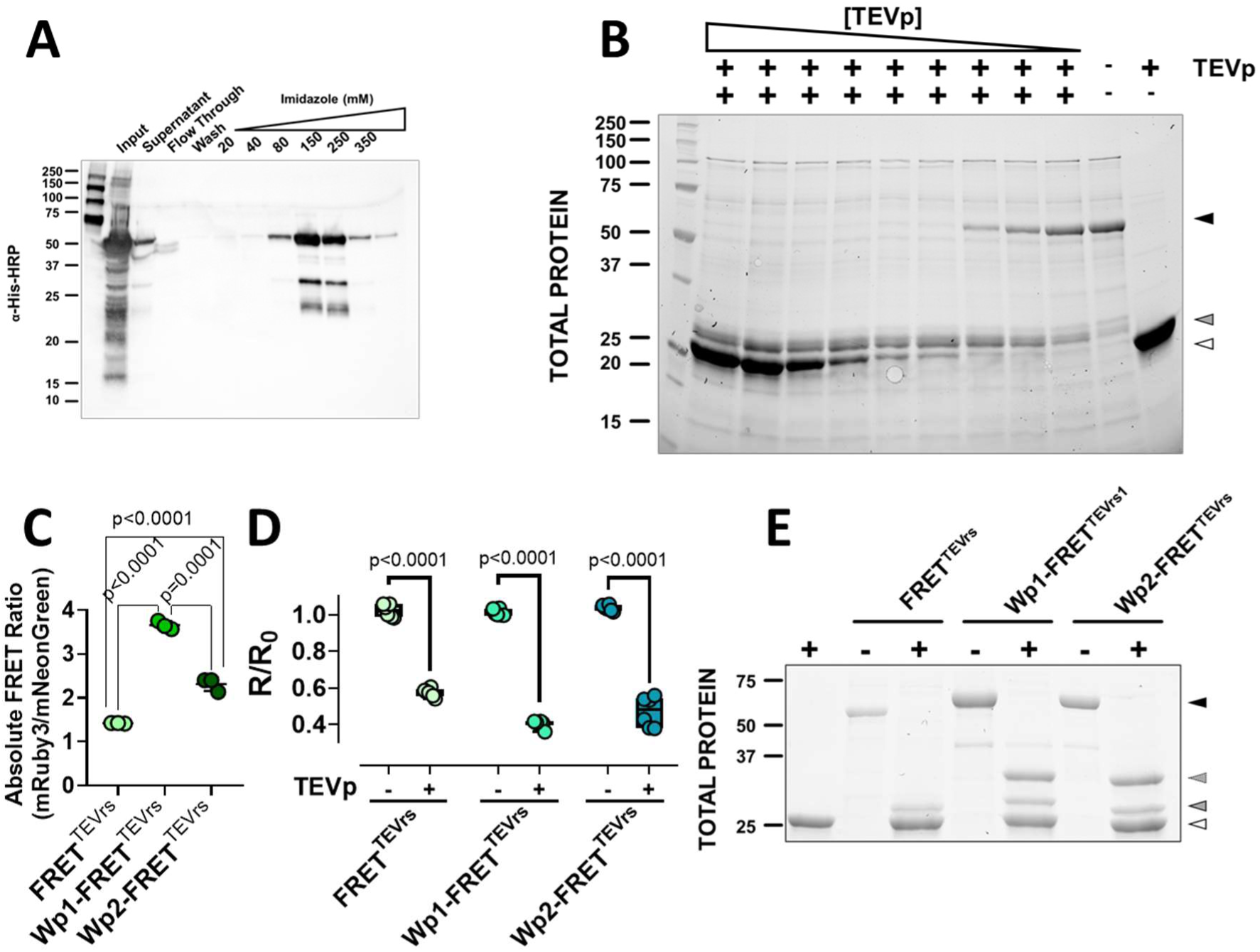
Recombinant FRET sensors are cleaved and report fluorescent changes in vitro. **A)** Recombinant purification of FRET^TEVrs^ in *E. coli* BL21. Different fractions were loaded at each lane as indicated in the upper panel. Elution was done at an increasing imidazole concentration. **B)** *In vitro* FRET^TEVrs^ incubation with TEVp at different concentrations show an increasing effect at higher concentration of TEVp. (Sizes marked on the right of the gel corresponding to dark arrow: FRET^TEVrs^ full size; grey arrow: processed FRET^TEVrs^; white arrow TEVp). **C)** Relative values of mRuby3/mNeonGreen of different FRET^TEVrs^ versions in absence of helpers and WW-Wp1 and WW-Wp1 combination (n=3 repeats, p-values are indicated according to ordinary one-way ANOVA). **D)** Normalized ration values (mRuby3/mNeonGreen) to ratio fluorescence of each FRET^TEVrs^ sensor before and after addition of TEVp (n=6 repeats, p-value indicated two-tailed nonparametric Mann– Whitney test). **E)** Gel displaying correct processing of the different FRET^TEVrs^ sensor in presence and absence of TEVp.

**Supplementary figure 2.**
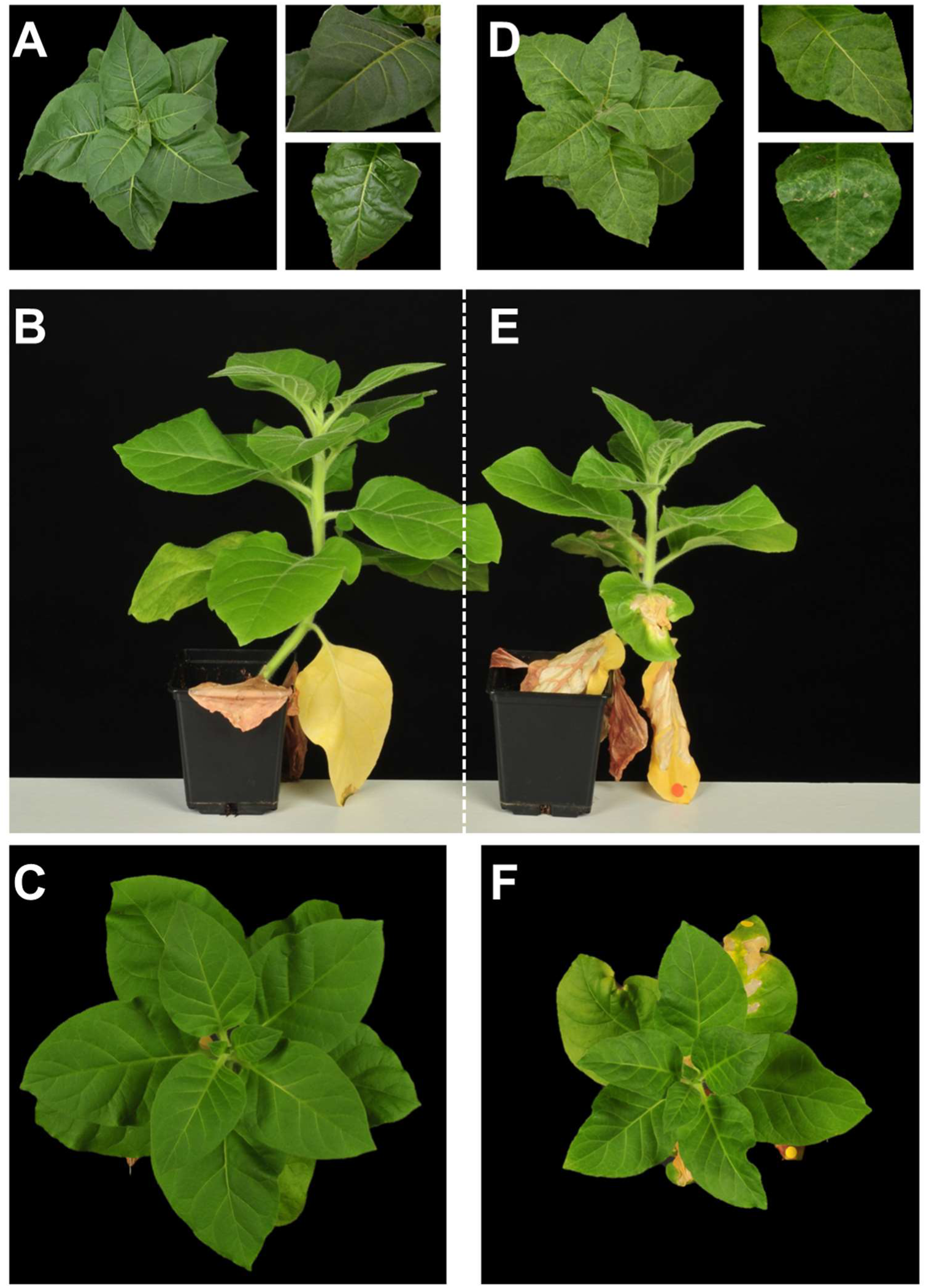
Impact of Tobacco-Etch Virus on of *N. tabacum* infected plants. Photography of tobacco plants used for assessing FRET^TEVrs^ performance during viral infection. **A-C)** Tobacco healthy plants. **D-F)** Nicotiana tabacum plants inoculated with TEV virus.

**Supplementary figure 3.**
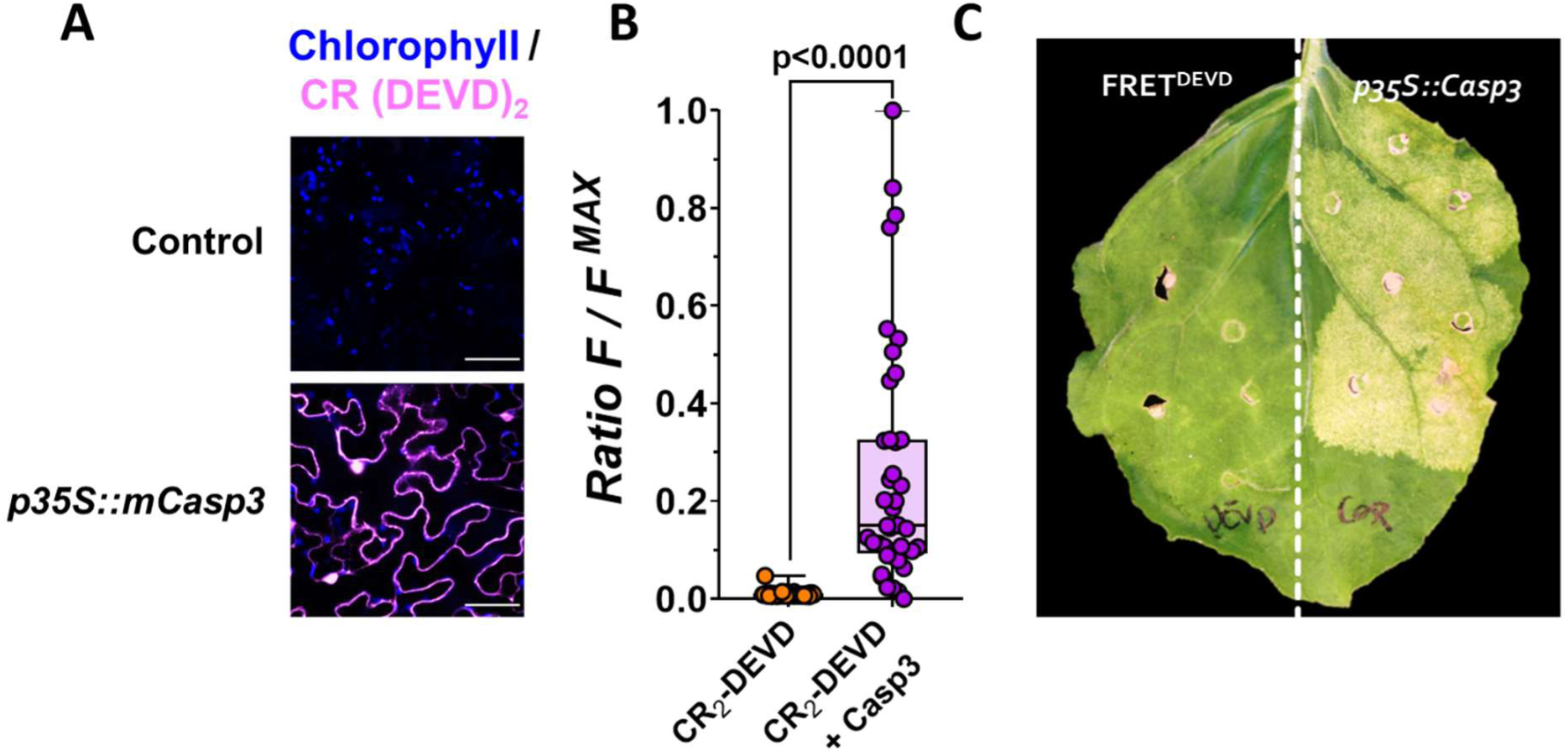
Caspase-3 is active in plants. **A)** Confocal images of *N. benthamiana* epidermal cells in leaves treated with CR2-(DEVD) after transient expression of caspase-3 under 35S promoter. CR2-(DEVD) is depicted in purple and chlorophyll autofluorescence is shown in blue. **B)** Dot-box showing normalized quantification of fluorescent values of CR2(DEVD) in absence and presence of capase-3 (orange and purple respectively; n≥41 biological repeats, t-test p-value<0.0001). **C)** Evident signs of cell-death phenotype observed in *N. benthamiana* induced by transient expression of caspase-3, at 6 days after infiltration.

**Supplementary figure 4.**
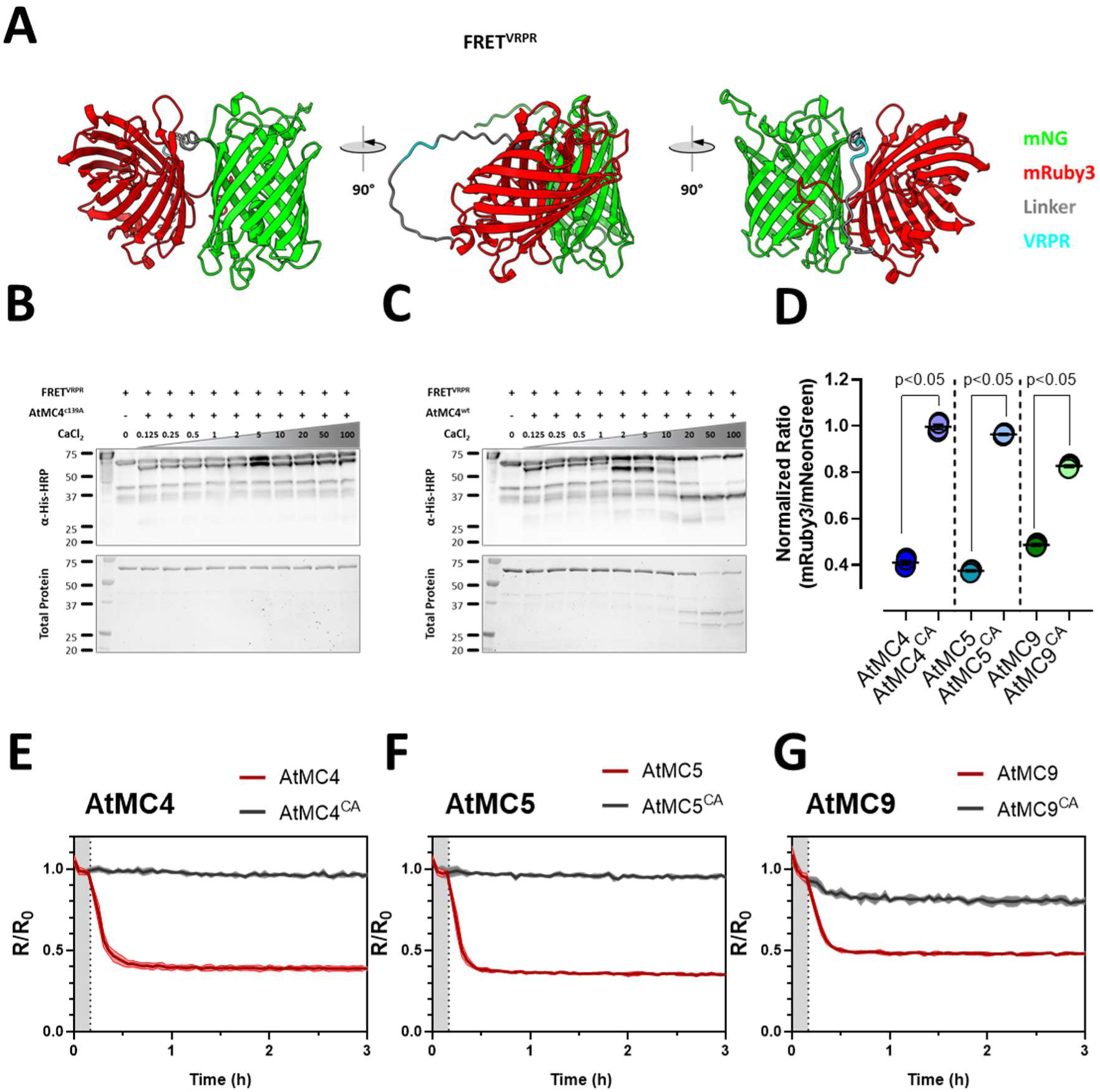
In vitro characterization of a customized metacaspase specific FRET sensor. **A)** Structural protein prediction using AlphaFold2 displaying the disposition of the FRET^VRPR^ sensor features from different angles. mNeonGreen (green), mRuby3 (red), and linker sequence (grey) remarking the minimal recognition site for metacaspases VRPR (blue). **B, C)** *In vitro* processing of FRET^VRPR^ sensor at increasing levels of calcium with co-incubated with inactive AtMC4C139A **(B)** and active AtMC4**(C)**. **D)** Normalized ratio values (mRuby3/mNeonGreen) of FRET^VRPR^ incubated *in intro* with active and inactive versions of AtMC4, AtMC5 and AtMC9 after 30 minutes in presence of 10mM CaCl2 (p-value <0.05 according to Mann-Whitney test, n=4 repeats). **E-G)** Time-dynamics response of the normalized ratio (mRuby3/mNeonGreen) of FRET^VRPR^ when incubated with active versions of AtMC4, AtMC5 and AtMC9 (red lines) or their inactive versions (grey line).

**Supplementary figure 5.**
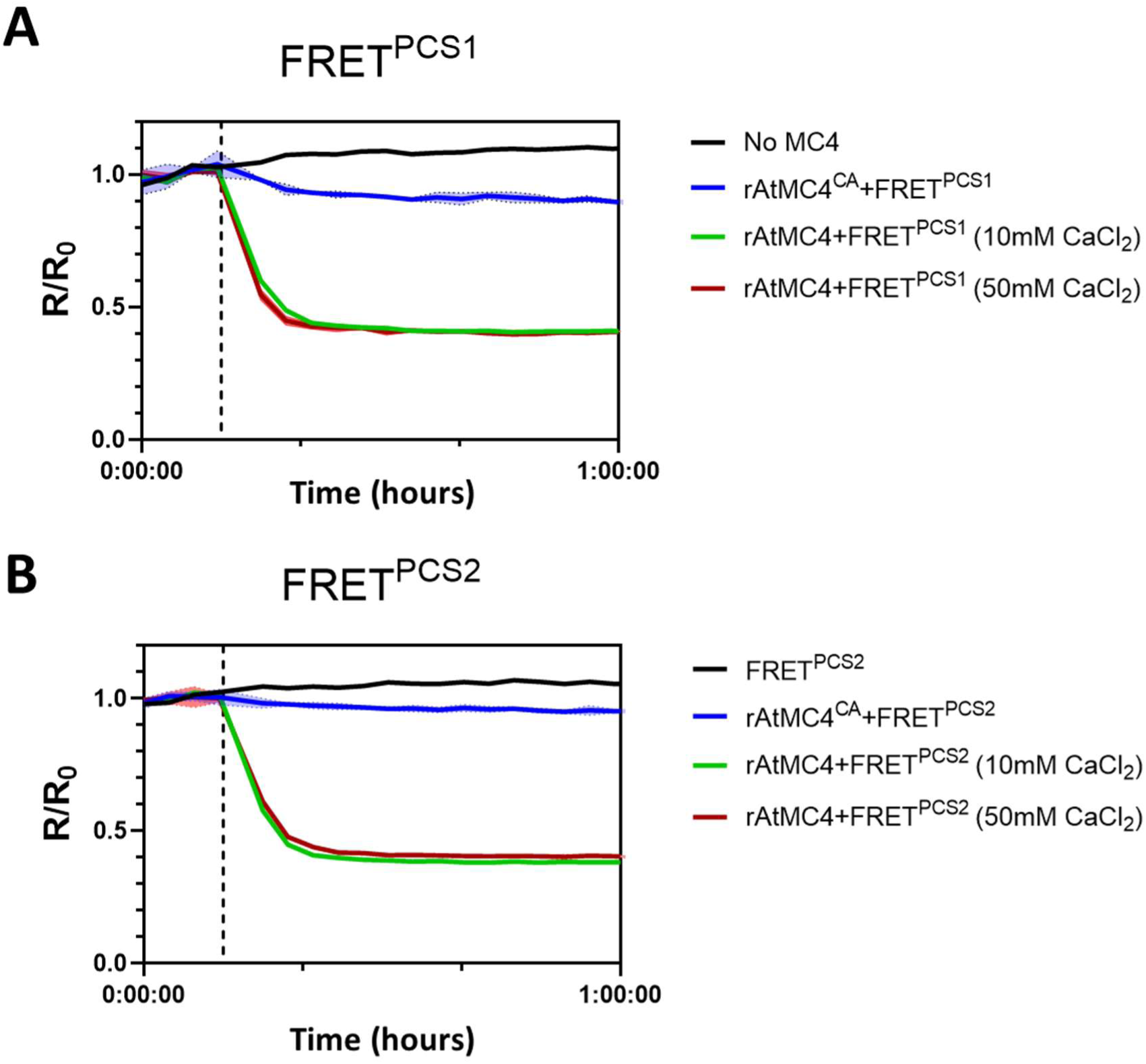
Set of FRET^PCS1^ and FRET^PCS2^ biosensors validation *in vitro*. **A-B)** Dynamics of normalized quantification of the FRET ratio (mRuby3/mNeonGreen) of recombinantly produced PROPEP1-based FRET biosensors FRET^PCS^^1^ and FRET^PCS^^2^ after CaCl2 addition indicated by vertical dotted arrow of mock conditions (buffer in black), incubated with inactive AtMC4 (rAtMC4^CA^; blue), and with active AtMC4 at 10 mM and 50 mM concentrations of CaCl2 (green and red, respectively).

**Supplementary figure 6.**
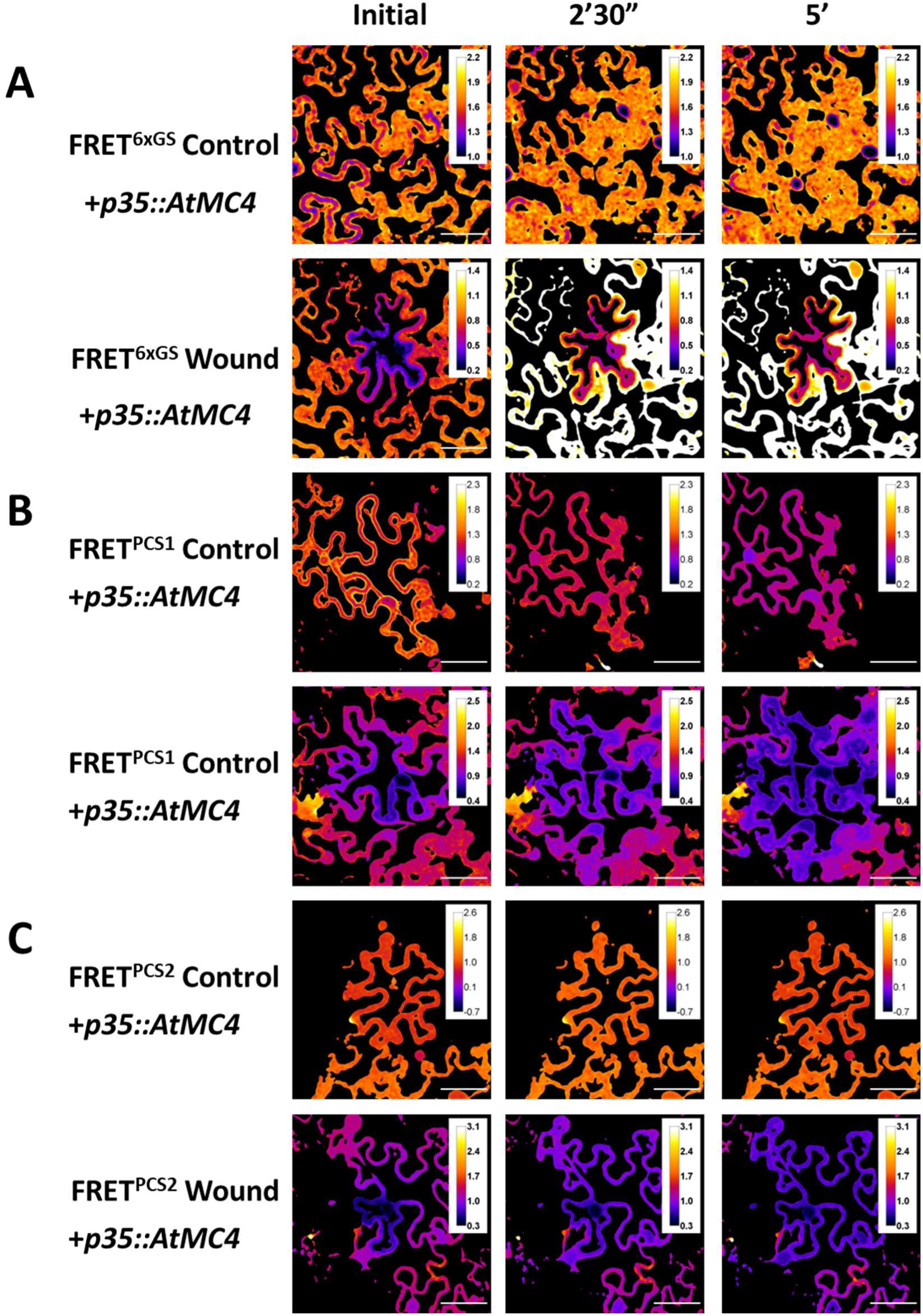
Ratio images of mRuby3/mNeonGreen of different versions of FRET reporters expressed in *N. benthamiana* in normal conditions (control) or undergoing laser ablation (wounding) at different timepoints. **A)** FRET^6xGS^. **B)** FRET^PCS1^. **C)** FRET^PCS2^. Leaves were co-infiltrated with their corresponding FRET sensors and a cassette overexpressing AtMC4 (pB7WG2,0-AtMC4s). Images were taken two days after infiltration. (Bars represent 50 μm).

**Supplementary figure 7.**
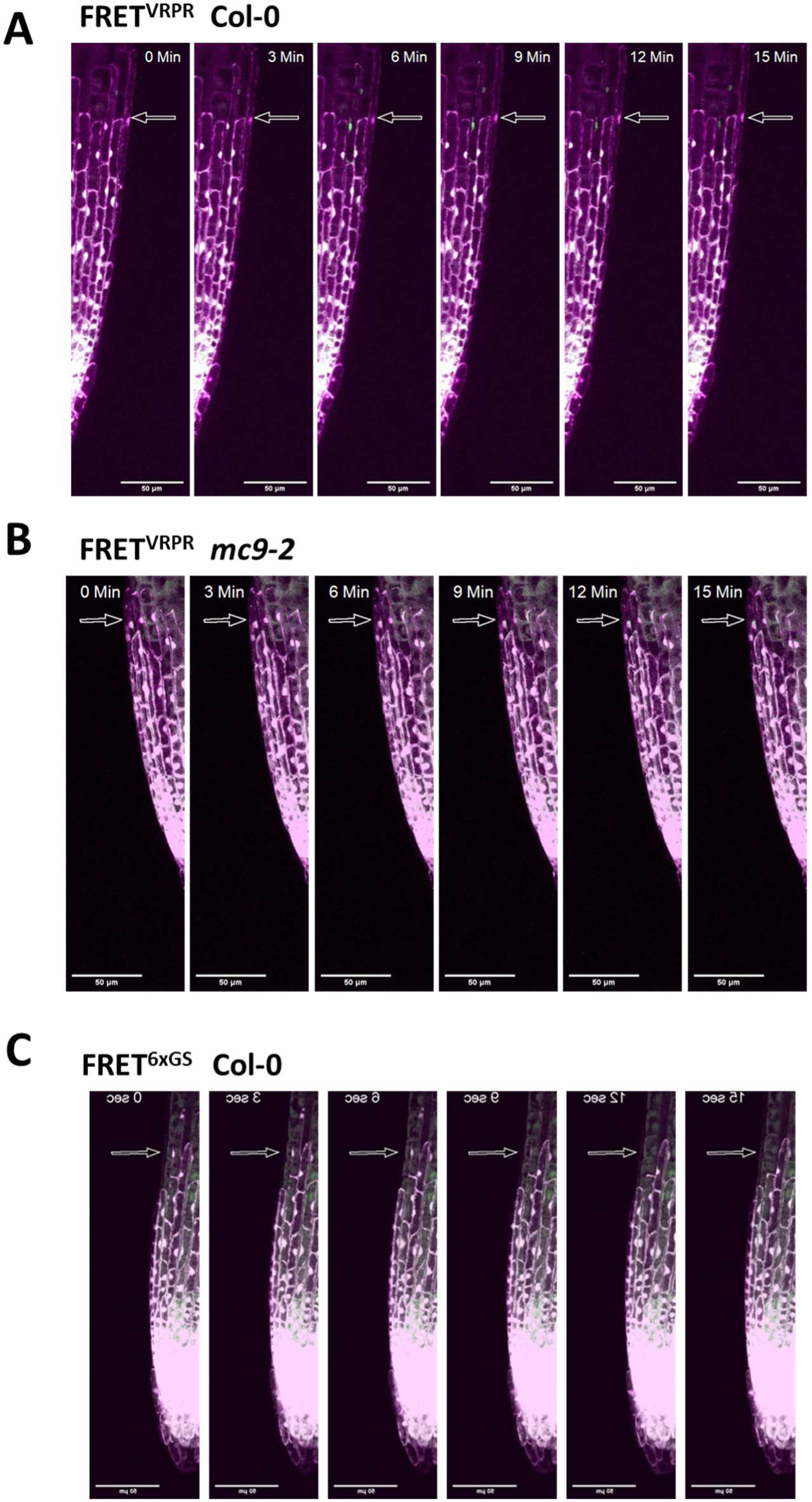
Confocal images of Arabidopsis roots expressing FRET biosensors reveal metacaspase activity in late events of lateral root cap. FRET^VRPR^ expressed in Col-0 (A)*, mc9-2* background **(B)** and FRET^6xGS^ in Col-0 **(C).**

**Supplementary video 1.** Timelapse video of the calculated FRET ratio signal of FRET^PCS1^ undergoing wounding

**Supplementary video 2.** Timelapse video of the calculated FRET ratio signal of FRET^6xGS^ in normal conditions.

